# Learning temporal structure engages hippocampus and guides value-based behaviour

**DOI:** 10.64898/2026.04.24.720548

**Authors:** Svenja Nierwetberg, David Orme, Maxwell Chen, Andrew F. MacAskill

## Abstract

The ability to learn and exploit structured relationships between events is fundamental to adaptive behaviour and episodic-like memory, yet the neural mechanisms that support such learning remain poorly understood. Progress has been limited by the difficulty of dissociating relational structure from sensory cues, value, and motor output in experimentally tractable tasks. Here we introduce a temporally structured olfactory task for mice that isolates relational structure from cue identity and reward statistics. Mice rapidly learned to use the ordered relationship between two sequentially presented odours to predict outcome. Because individual odours were equally associated with reward across contexts, their predictive value depended entirely on their relationship to preceding cues. Behavioural analyses and reinforcement learning models revealed a gradual shift to the use of structured representations as learning progressed. Dopamine activity in the nucleus accumbens reflected this transition. Over learning, identical sensory inputs became associated with opposite predicted outcomes depending only on their temporal position, consistent with dopamine prediction errors reflecting latent task structure rather than cue identity. Trial-by-trial variability in dopamine dynamics was quantitatively captured by the same models that described behavioural strategy, linking dopamine prediction errors directly to these learned structural representations. Finally, using a matched control task that preserved sensory input and reward statistics while eliminating structure, we found that learning and using temporal structure selectively increased ventral hippocampal activity. Together, these results establish a minimal, non-spatial paradigm for studying structural learning in mice, link behavioural representations to dopamine signalling, and show that isolating relational structure from sensory and reward confounds selectively recruits hippocampal circuitry long theorised to support it.

## INTRODUCTION

Learning and exploiting relationships between stimuli is fundamental to adaptive behaviour^1–3^. In many situations, the meaning of a cue cannot be inferred in isolation, but depends on its relationship to other events in space or time^1,4,5^. More generally, the ability to represent and use structured relationships between distinct cues underlies the associations that form the basis of episodic and semantic memory^6,7^.

Consistent with this central role, impairments in memory-guided behaviour that depend on structured relationships are a hallmark of a wide range of neurological and psychiatric disorders, including Alzheimer’s disease, schizophrenia, depression, and anxiety disorders^8–11^. Despite this, progress has been limited by the difficulty of isolating relational structure from sensory cues, reward statistics, and motor output in experimentally tractable tasks^12–15^. While much is known about how individual cues become associated with outcomes, how sequences of cues that share overlapping elements are combined to predict distinct outcomes remains unclear.

A prominent proposal is that such learning depends on the medial temporal lobe, and in particular the hippocampal formation^1,7,12^. Hippocampal activity is strongly modulated by temporal and spatial context, is sensitive to the order in which events are experienced, and is required for differentiating appropriate behaviour towards identical cues encountered in different contexts^4,16–23^. These properties suggest a role in representing relationships between events rather than their sensory features alone.

Most experimental investigations of these ideas have focused on spatial navigation, where structure is inferred from movement through space^3,7,16,24^. While these approaches have provided important insights, their interpretation is often complicated by continuous variables and alternative solution strategies that do not require explicit representation of relational structure. Moreover, the principles of structural learning extend beyond spatial cognition^1,2,19,25–29^, and many disease-associated deficits are not primarily spatial.

Outside the spatial domain, progress has been limited by the lack of behavioural paradigms that isolate relational structure while remaining experimentally tractable^12–14,30^. Many non-spatial tasks admit alternative solutions based on elemental or configural strategies, making it difficult to determine when behaviour truly depends on relationships between cues^13^. As a result, it remains challenging to assign a specific role to hippocampal circuitry or to relate behaviour to underlying neural signals.

Here we address this gap by introducing a temporally structured olfactory task that isolates relational structure from cue identity and reward statistics. By presenting cues sequentially and counterbalancing both cue identity and outcome, the task ensures that individual cues are uninformative and that predictive value depends entirely on their relationship across time. Efficient performance therefore requires disambiguating overlapping cue sequences based on temporal context, providing a minimal and experimentally tractable assay of relational structure and enabling direct tests of when hippocampal circuitry is engaged by such representations.

Using this paradigm, we show that behaviour transitions from cue-based to relational representations across learning. Dopamine release in the nucleus accumbens provides an independent readout of value learning^27,31–33^, demonstrating that identical sensory inputs come to predict different outcomes depending on their temporal context. Finally, using a matched control task that preserves sensory input and reward statistics while eliminating relational structure, we show that learning and using temporal structure selectively engages ventral hippocampus.

Together, these results establish a minimal, non-spatial paradigm for studying structural learning in mice, demonstrate that animals use relational structure to guide value-based predictions, and identify hippocampal circuitry as selectively recruited when such representations are required.

## RESULTS

### A temporally structured olfactory task that isolates relational cue information

We designed a structural learning task (Fig. 1A-C), adapted from classic biconditional discrimination and spatial navigation studies^13,20,29,34–36^, with the goal of creating a behavioural paradigm in which the appropriate response on each trial depends solely on the relationship between distinct cues.

**Figure 1.**
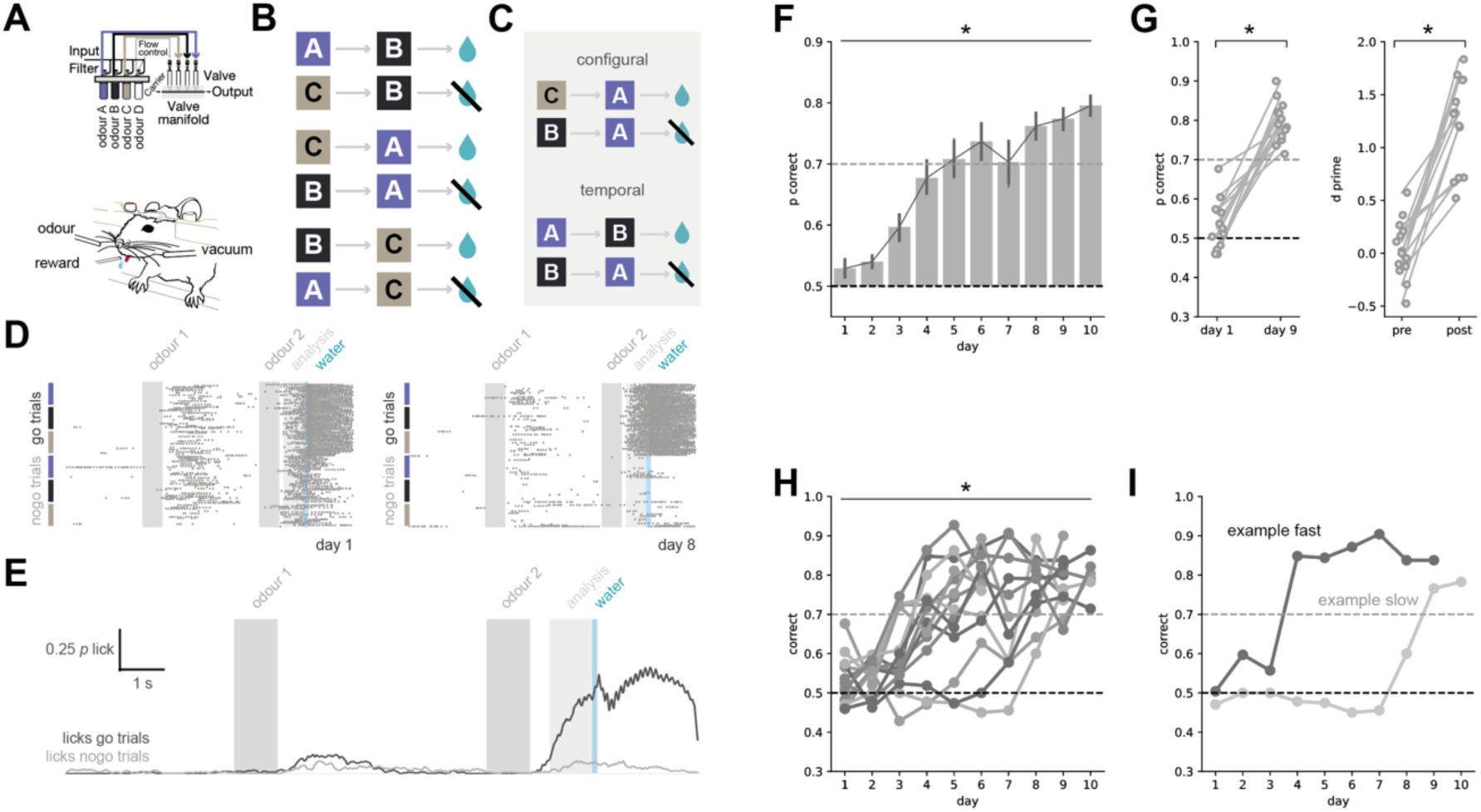
Mice rapidly learn a structured paired-associates task. **A.** Schematic of the behavioural setup. Odours were delivered to the mouse’s nose via pressurised solenoid valves and cleared by a constant vacuum stream. Reward was delivered through a metal lick spout. **B–C.** Task design. Three odours were presented in counterbalanced pairs such that each odour appeared equally often in the first and second position and was equally associated with rewarded and unrewarded outcomes. Accurate performance therefore requires learning of task structure, combining configural information (the meaning of an odour depends on its paired partner) and temporal order (e.g. A→B is distinct from B→A), as illustrated in **C**. **D.** Example lick rasters from a single novice session (left) and expert session (right). In both go and no-go trials, the second odour is identical (colour-coded at left). Despite this, expert mice selectively lick in anticipation of reward on go trials but not on no-go trials. **E.** Mean lick rate across mice for go (black) and no-go (grey) trials in expert animals. **F.** Proportion of correct trials across training days. **G.** Performance on the first and last training days for each mouse, shown as proportion correct (left) and d′ (right). **H.** As in **F**, shown for individual mice to illustrate variability in learning rate. **I.** Example learning curves from a representative fast-learning mouse and a slow-learning mouse.

Classically, this has been achieved by designing tasks in which the signals for reward (+) and no reward (–) are composed of the same elements, but differ in how those elements are combined. A commonly used example is a biconditional discrimination, in which pairs of four cues define trial outcome (for example A|B+, A|Y–, X|Y+, X|B–). In such designs, individual elements (A, B, X or Y) are uninformative, and correct performance requires sensitivity to cue combinations.

However, an important limitation of these tasks is that they can be solved using multiple strategies. While they can be solved using relational or structural learning, they can also be solved by binding cues into a single percept, such that ‘AB’ is treated as a distinct sensory stimulus from ‘AY’. This ambiguity has contributed to inconsistent results in the literature, particularly when attempting to assign a specific role to hippocampal circuitry.

To overcome this limitation, we took inspiration from a series of spatial navigation studies and introduced two key modifications to the classical design^12,37^. First, we separated each cue in time, emphasising their discrete nature and reducing perceptual binding. Cues were therefore presented sequentially (for example A followed by B), rather than simultaneously (A|B). Second, we counterbalanced cue pairs such that the same pair could be either rewarded or unrewarded across the task depending on the order they were presented in.

Specifically, we used three odours (A, B and C), arranged into six possible ordered pairs (A→B +, B→A -, C→A +, A→C -, B→C +, C→B -). These pairs were reinforced such that correct performance required differentiating both the identity of the odour pair and the order in which cues were presented (for example B after A is rewarded, whereas A after B is not). Under this design, each odour appears equally often in rewarded and unrewarded contexts, and each unordered pair of odours is both rewarded and unrewarded (Fig. 1B). As a result, simple elemental or configural representations are not sufficient: predictive value depends entirely on the temporal relationship between cues (Fig. 1C). Efficient performance therefore requires disambiguating overlapping cue sequences based on temporal context.

Using this design, water-restricted, head-fixed mice were presented with randomly interleaved rewarded and non-rewarded odour pairs. On each trial, the first odour was delivered for 1 s on a constant carrier stream, followed by a 5 s delay, before presentation of the second odour for 1 s. On rewarded (‘go’) trials, a water reward was delivered following a 1.5 s trace interval after the second odour. Importantly, reward delivery was independent of licking behaviour, such that all go trials were followed by reward in a Pavlovian manner. Licking was recorded continuously using a capacitance-based lick sensor. Learning was quantified as the emergence of anticipatory licking during the trace interval on go trials, and the suppression of anticipatory licking on no-go trials.

### Mice rapidly acquire the structural learning task

Before training on the full task, mice were habituated to head fixation and to the overall structure of odour delivery. Habituation consisted of 2–3 days of head fixation for progressively increasing durations, starting from 1–2 minutes and increasing to 10 minutes. During these sessions, mice received random water rewards from the lick spout to encourage licking behaviour.

Mice were then trained on a simplified version of the task containing only rewarded (‘go’) odour combinations (A→B, B→C and C→A). During this stage, mice learned to exhibit anticipatory licking for water reward during 30-minute daily sessions. Mice were advanced to the full task once they reliably showed anticipatory licking on at least 80% of trials across two consecutive days. On average, mice required ∼7 days to complete this training stage.

On the first day of the full task (day 1), mice behaved suboptimally. Anticipatory licking was observed on both rewarded (‘go’) and unrewarded (‘no-go’) trials, and substantial licking was also evident following presentation of the first odour in each pair (Fig. 1D). As a result, overall task performance on day 1 was close to chance.

However, performance improved rapidly across subsequent training days. Within 8–10 days, all mice exhibited high and stable performance, characterised by robust anticipatory licking selectively on go trials, minimal licking on no-go trials, and a marked reduction in licking following the first odour (Fig. 1D,E). We quantified this learning in two complementary ways (Fig 1G). First, the proportion of correct trials increased significantly between the first and final training sessions. Second, behavioural sensitivity, quantified using the signal detection theory metric *d′* (reflecting the ability to predict trial type based solely on licking behaviour), also increased markedly across training.

Although all mice reached greater than 70% correct performance by the end of the training period, learning rates varied substantially across individuals (Fig. 1H,I). Some mice acquired the task within only a few days, whereas others required more than a week to reach criterion performance.

Together, these results demonstrate that mice can rapidly learn to use the temporal structure of olfactory cues to guide behaviour in this task.

### Mice rely on discrete olfactory cues rather than non-olfactory signals or residual odour

Olfactory tasks can be confounded by non-olfactory cues associated with stimulus delivery, such as valve sounds or airflow changes, as well as by residual odour persisting across time^38,39^. Although our olfactory delivery system provides precise temporal control^40^, we explicitly tested whether mice were using odour identity in the manner intended by the task.

In well-trained mice, we first performed control sessions in which airflow, odour identity, or both were removed, leaving only auditory valve clicks and/or pressure changes (Fig. 1 - Figure Supplement 1). Under these conditions, performance collapsed to chance and recovered immediately when standard odour delivery was reinstated, indicating that behaviour depends on odour identity rather than non-olfactory cues.

We next asked whether residual odour during the delay period could allow mice to encode cue pairs as a single continuous percept. Photoionisation detection measurements confirmed that odour concentration decayed rapidly following cue offset, reaching levels orders of magnitude lower than those during cue presentation by the end of the delay (Fig. 1 - Figure Supplement 1). To test whether such residual signals could nevertheless support performance, we introduced probe trials in which a low concentration of odour was delivered continuously throughout the delay period.

Critically, the concentration of this probe odour was calibrated to exceed residual levels measured in standard trials, while remaining substantially lower than the concentration during normal odour presentation. This manipulation therefore provided a sustained olfactory signal during the delay that was stronger than any residual odour available under standard conditions.

Under these conditions, performance on probe trials dropped markedly, despite preserved performance on interleaved standard trials. Thus, providing a continuous low-level odour signal during the delay does not support task performance, even when this signal exceeds natural residual levels.

Together, these results indicate that mice solve the task by integrating information from two discrete olfactory cues separated in time, rather than by exploiting continuous sensory signals or non-olfactory cues.

### Behavioural choices increasingly reflect ordered cue relationships across learning

To quantify which aspects of the task guided behaviour, we fit a logistic regression model to trial-by-trial choices that included predictors corresponding to different interpretations of cue identity (Fig. 2A, B). Elemental predictors indicated the presence of individual odours, configural predictors indicated unordered cue pairs, and structural predictors indicated specific odour pairs in a defined temporal order. Predictors reflecting reward and choice on the previous trial were included to capture the influence of recent trial history.

**Figure 2.**
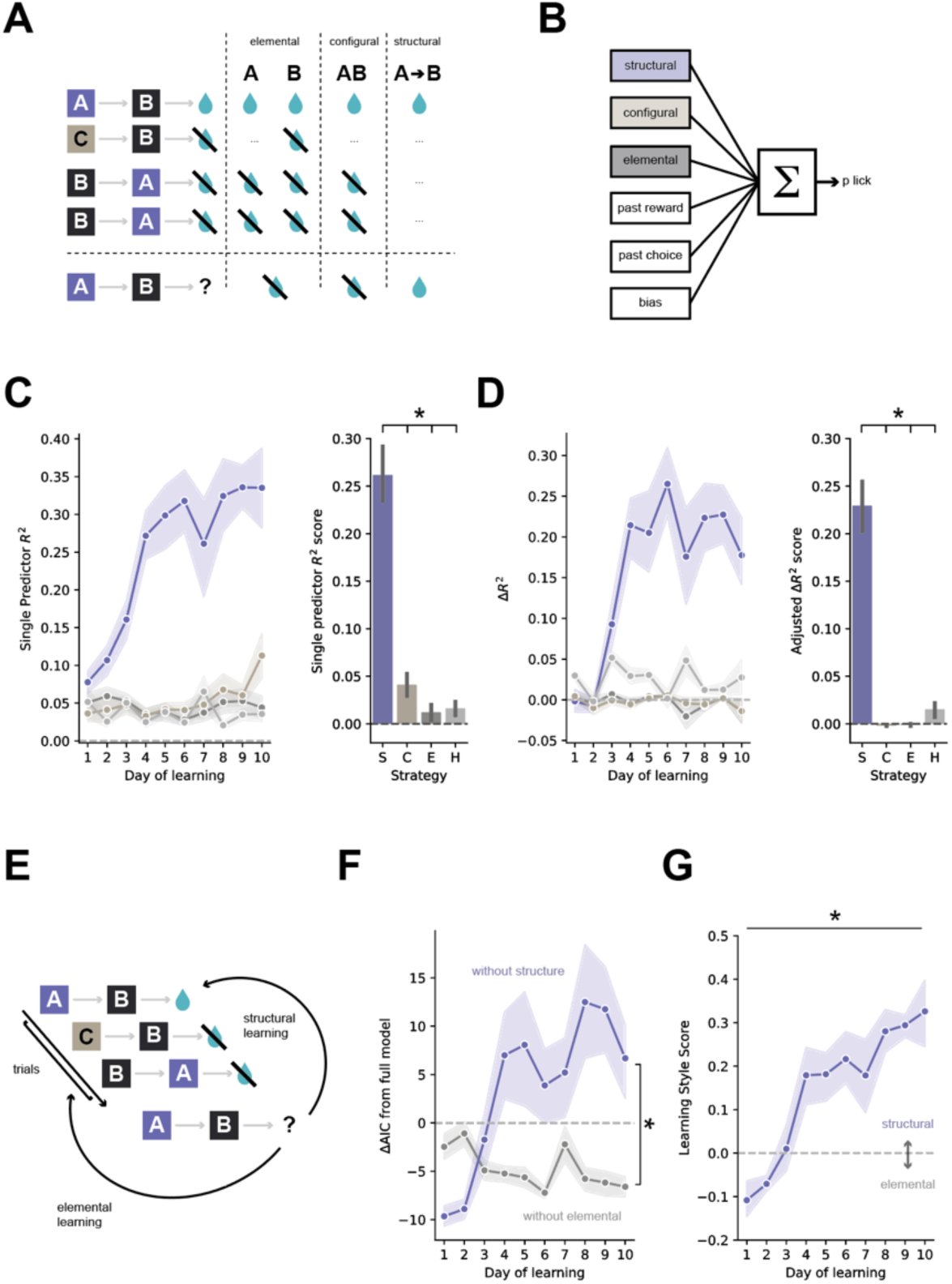
Mice progressively adopt structure-based strategies across learning. **A.** Schematic illustrating how elemental, configural, and structural strategies generate distinct predictions across a sequence of trials. Elemental strategies assign value to individual odours, configural strategies to specific odour pairs, and structural strategies assign value according to ordered cue relationships. **B.** Regression framework used to predict trial-by-trial licking behaviour. Predictors captured elemental, configural, and structural information, along with trial and reward history (see Methods). **C.** Pseudo-R² across learning for individual predictor classes: structural (purple), configural (brown), elemental (dark grey), and history (light grey). Structural predictors explain an increasing proportion of behavioural variance over training (left) and dominate in expert animals (right). **D.** Unique variance explained by each predictor class, quantified as the change in pseudo-R² following removal from the full model. Structural predictors account for the largest unique contribution in expert animals. **E.** Schematic of reinforcement learning models used to capture behaviour. Elemental models update values assigned to individual odours, whereas structural models update values over shared relational representations within a Rescorla–Wagner framework. **F.** Model comparison using ΔAIC relative to the full model. Removing structural updates (purple) produces a larger decrease in model fit than removing elemental updates (grey), indicating a greater contribution of structural learning. **G.** Learning style score, defined as the difference between structural and elemental model fits across training. Positive values indicate structural dominance. Mice show a gradual shift from reliance on elemental to structural strategies over learning.

We first examined regression coefficients from the full, cross-validated model (Fig. 2 – Figure Supplement 1). Across mice, predictors corresponding to rewarded structural sequences had large positive weights, while those corresponding to unrewarded sequences had large negative weights, indicating a strong influence on choice behaviour. In contrast, coefficients associated with elemental, configural, and trial-history predictors were near zero.

Importantly, this structure dependence was not static. When regression models were fit separately to each training session, the contribution of structural predictors increased systematically across learning. Structural predictors dominated late in training, where reduced models containing only structural predictors increasingly outperformed models containing elemental, configural, or history-based predictors as training progressed

(Fig. 2C). Similarly, shuffling structural predictors in the full model produced a progressively larger reduction in model performance across days, whereas shuffling other predictor classes had little effect at any stage of learning (Fig. 2D).

Together, these analyses indicate that behavioural control shifts over training towards reliance on ordered cue relationships, as revealed by a time-resolved regression analysis that makes no assumptions about the underlying learning rule.

### Value assignment increasingly reflects structural representations across learning

To examine how learning strategy evolves across training, we implemented a series of reinforcement learning models based on a simple Rescorla–Wagner learning rule^29^. Across all models, value estimates were updated proportionally to the prediction error on each trial, ensuring that differences in model behaviour reflected differences in how sensory cues were represented, rather than differences in the learning rule itself.

In the task, mice make a binary choice (‘go’ or ‘no-go’), but the predicted value of each choice depends on how the odour cues are interpreted. In an elemental representation, each odour contributes independently to the predicted value of a choice. In a structural representation, the value of a choice depends on the specific ordered pair of odours presented on that trial. We therefore defined separate state spaces corresponding to elemental and structural interpretations of the task (Fig. 2E). Formally, the structural model can be interpreted as a discrete latent-state representation, in which each ordered cue pair defines a distinct hidden state. Under this view, learning corresponds to assigning value to inferred task states defined by cue relationships, rather than to observable cues themselves.

We fit these models independently to trial-by-trial choices from each mouse and each training session. To allow for the possibility that behaviour reflected a mixture of strategies, we also fit a hybrid model in which predicted choice values were computed as a weighted combination of elemental and structural value estimates, with separate inverse temperature parameters governing the influence of each representation on choice (see methods).

Across the majority of sessions, models incorporating a structural state representation provided better fits to behaviour than models based on elemental representations alone or a random-choice baseline (Fig. 2F). However, the hybrid model consistently outperformed a purely structural model, despite the additional penalty for increased model complexity. Consistent with this, in very early sessions, models incorporating an elemental state representation consistently outperformed those based on structure. This indicates that although structural representations dominated behaviour, elemental representations also contributed to choice, especially in early training.

To summarise this transition, we computed a structural–elemental index based on the relative inverse temperature parameters associated with each representation (Fig. 2G). This index increased systematically across training days, indicating a gradual shift from elemental to structural control of behaviour. Importantly, model fits on early training days were substantially better than chance, indicating that this transition reflects a change in representation rather than poor model performance early in learning.

Together, these results indicate that mice initially assign value based on individual cue identity, but progressively adopt a representation in which value is assigned to structured cue sequences as learning proceeds.

### Dopamine prediction error reflects learned value over task structure

If behaviour is guided by relational representations rather than individual cues, then value should depend on inferred task states defined by temporal context. We therefore asked whether dopamine prediction errors provide an independent readout of these learned representations^27,31,33^. To test this, we recorded dopamine activity in the nucleus accumbens during task performance across learning using the dopamine sensor dLIght^41^. We used this to compare early sessions, when behaviour was near chance, to expert sessions in which mice reliably used cue order to predict outcome.

We first examined trial-averaged dopamine activity aligned to cue presentation and reward delivery on rewarded (‘go’) and unrewarded (‘no-go’) trials (Fig. 3A). In early training sessions, dopamine responses were dominated by a robust response to reward delivery, with little differentiation between go and no-go trials during the interval between cue presentation and outcome. In expert sessions, dopamine activity increased selectively during the interval between the second odour and reward on go trials, while responses following reward delivery were markedly reduced.

**Figure 3.**
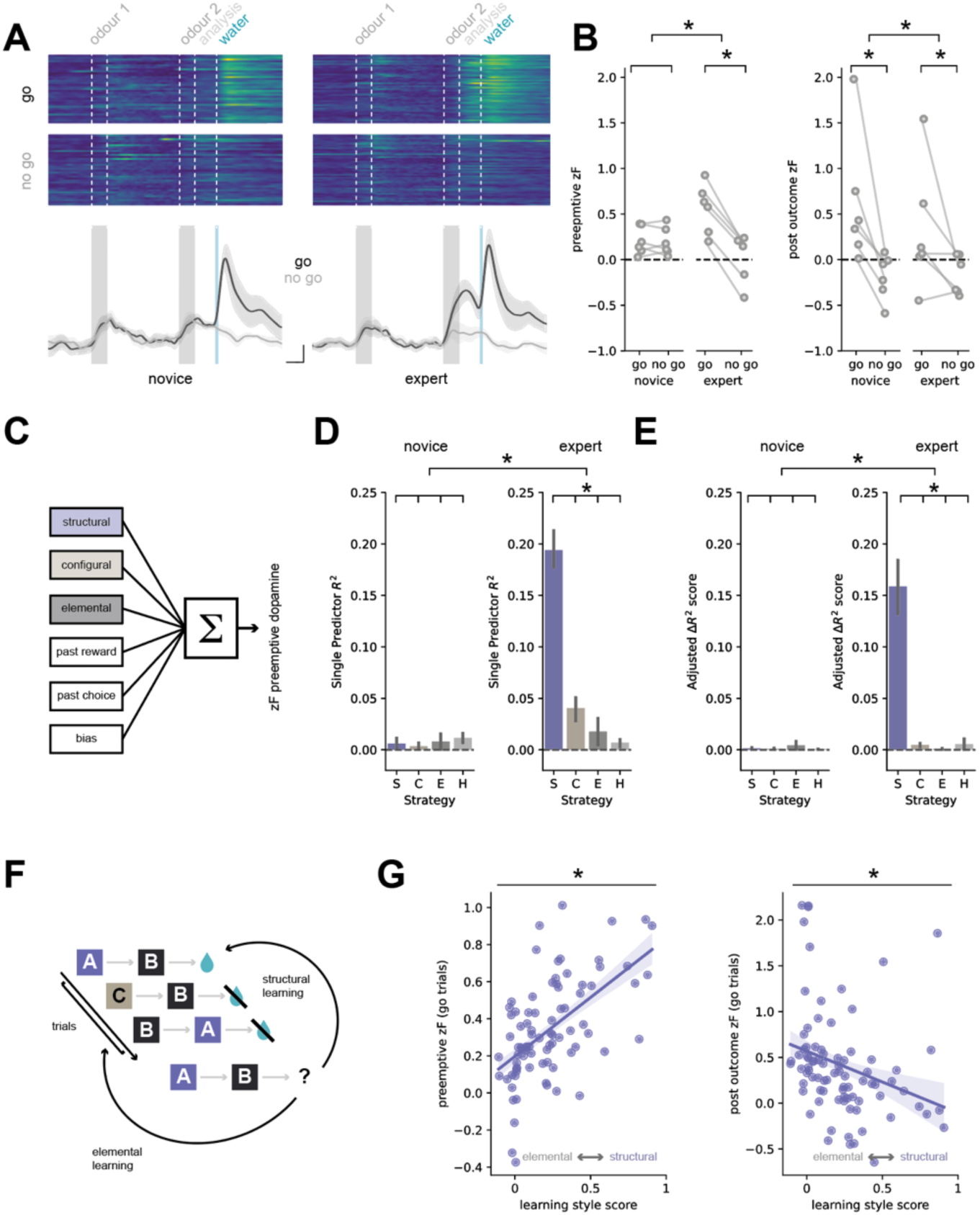
Dopamine signals reflect learned task structure. **A.** Top, trial-by-trial heatmaps of dopamine activity for go and no-go trials in novice (left) and expert (right) mice. Bottom, mean dopamine traces across mice for go (black) and no-go (grey) trials. In expert animals, dopamine responses diverge following the second odour, reflecting predictive encoding of trial outcome prior to reward delivery. Scale bars = 1 s, 0.5 zF. **B.** Quantification of dopamine signals in novice and expert mice before outcome delivery (pre-emptive; left) and after outcome (right). In expert animals, pre-emptive dopamine activity differentiates go and no-go trials, consistent with learned predictions, while post-outcome responses reflect outcome-dependent updates. **C.** Regression framework used to predict pre-emptive dopamine activity (defined as the period between the second odour and outcome delivery), using the same predictor set as in Figure 2: structural, configural, elemental, and history variables (see Methods). **D.** Single predictor fits reveal no consistent contributions from any predictor class in novice animals, indicating an absence of structured dopamine signals early in learning, but reveals a large contribution of structural predictors to pre-emptive dopamine activity. **E.** As in D but for unique variance (Δpseudo-R²) In expert animals, unique variance analyses reveal a dominant contribution of structural predictors to pre-emptive dopamine activity, mirroring the behavioural shift towards structure-based strategies. **F.** Schematic of the reinforcement learning model used to capture behaviour and generate trial-by-trial estimates of learning variables. **G.** Relationship between behavioural learning style and dopamine activity. The learning style score (derived from model fits; Figure 2) correlates with both pre-emptive and post-outcome dopamine signals across sessions, indicating that dopamine reflects value updates structured by latent task representations.

This redistribution of dopamine signalling from outcome to predictive period matches classical descriptions of reward prediction error signalling. Critically, at the time points analysed, sensory input was identical across conditions: in both go and no-go trials, the same odours were presented, differing only in their temporal order (Fig. 3 – Figure Supplement 1). The emergence of differential dopamine activity during the predictive period therefore cannot be explained by cue identity, but is instead consistent with prediction error computed over the learned relationship between cues. Identical sensory inputs are associated with opposite predicted outcomes depending only on their relationship to the preceding cue, indicating that value assignment depends on relational context rather than cue identity.

We quantified these effects by measuring dopamine activity during the pre-reward interval and following reward delivery (Fig. 3B). Across mice, pre-reward dopamine activity increased selectively on go trials across learning, whereas post-reward dopamine responses decreased correspondingly. Together, these changes indicate that dopamine prediction errors progressively align with learned task structure.

We next asked whether dopamine responses, like licking behaviour, could be described using the same structural framework described above. To address this, we applied the regression model used to predict choice behaviour, but instead used it to predict trial-by-trial dopamine activity during the pre-reward interval (Fig. 3C-E).

Early in training, dopamine responses were weakly predicted by multiple classes of predictors, consistent with the absence of a stable task representation. In contrast, in expert sessions dopamine activity was strongly predicted by structural predictors corresponding to ordered cue relationships, while elemental, configural, and trial-history predictors contributed little explanatory power. This mirrors the pattern observed for behavioural choices, indicating that dopamine dynamics track the same learned structural representations that guide behaviour. These analyses demonstrate that dopamine activity becomes selectively aligned with task structure as learning proceeds.

Finally, we asked whether individual differences in learning strategy were reflected in dopamine dynamics. Using the reinforcement learning models we used to describe behaviour above, we quantified for each session the relative contribution of elemental and structural representations to behaviour. We then related this learning score to dopamine activity measured in the same sessions (Fig. 3F,G).

Across mice and sessions, greater reliance on structural representations was associated with higher pre-reward dopamine activity on go trials, consistent with stronger predictive signalling. Conversely, reliance on structural representations was negatively correlated with dopamine responses following reward delivery, consistent with reduced prediction error at outcome. These relationships were observed both across training days within individual mice and across mice at matched stages of learning.

Thus, variability in how mice represent task structure is reflected in dopamine dynamics, linking behavioural strategy and neuromodulatory signalling within a common framework. Together, these results show that dopamine provides a trial-by-trial readout of structure-dependent value during task performance.

### Learning a structured task selectively engages ventral hippocampus

A central challenge in linking behaviour to circuit-level recruitment is dissociating neural activity related to sensory stimulation, reward delivery, and motor output from activity related to the underlying computation. This is particularly problematic in tasks involving learning, where changes in performance are often accompanied by changes in motivation, reward expectation, and action selection. To address this, we designed a control task that was carefully matched to the structured task in sensory input, reward statistics, and motor demands, but eliminated the possibility of using structure to predict outcome (Fig. 4A).

**Figure 4.**
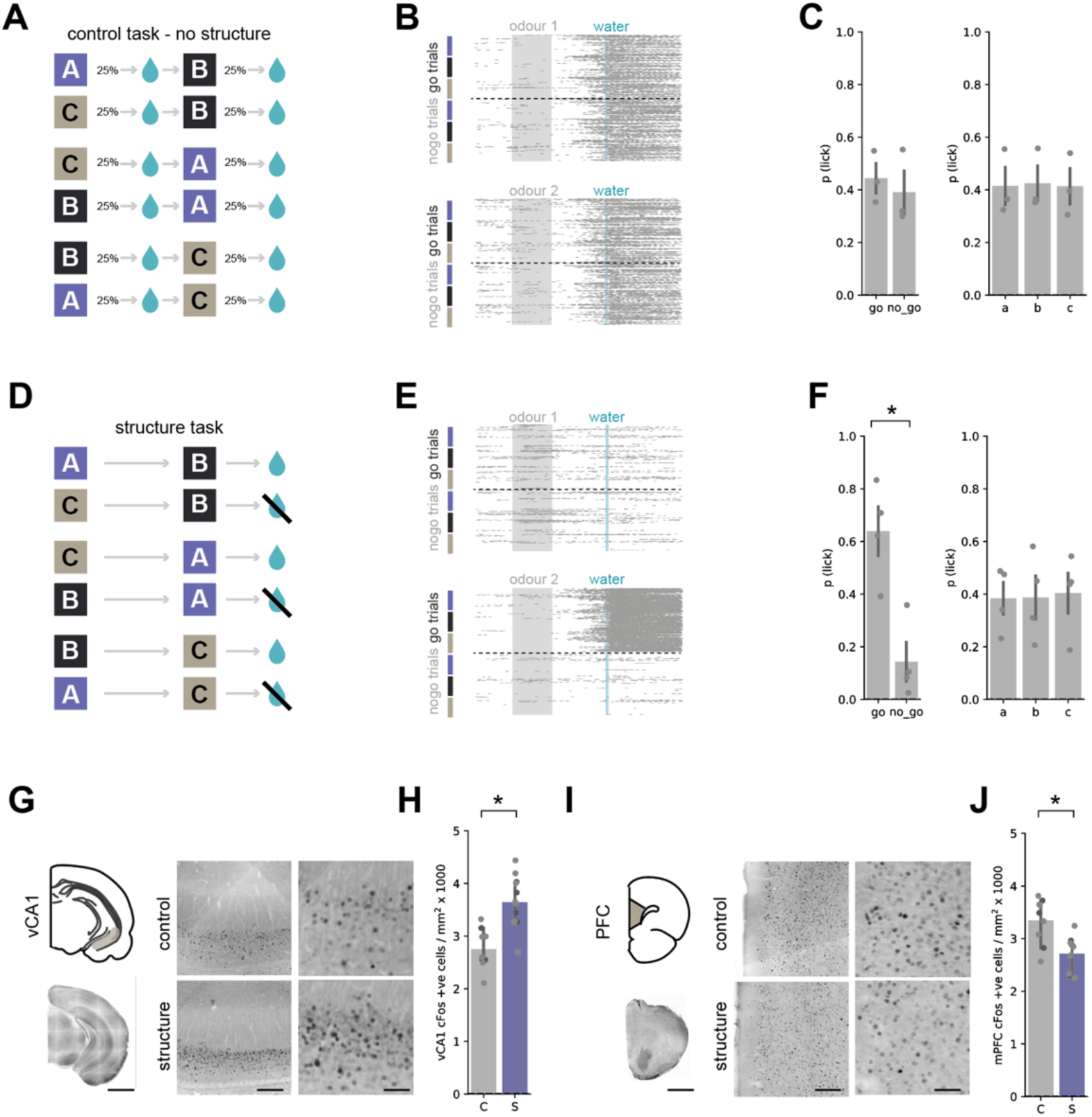
Isolating relational structure recruits ventral CA1. **A.** Design of the control task. Each odour was rewarded independently with a fixed probability (25%) on each presentation. This matches the structured task in terms of sensory input, reward frequency, and motor demands, but eliminates any predictive value of relational structure. **B.** Example lick rasters aligned to first and second odour presentations in the control task. Licking is equivalent across go and no-go trial types and across odour positions, indicating the absence of structure-based prediction. **C.** Left, mean lick probability to second odours corresponding to go and no-go trials in the control task. Right, mean lick rate to each individual odour, independent of position or trial type. Licking is uniform across conditions, confirming that behaviour cannot exploit structure. **D–F.** As in **A–C**, for mice performing the structured task. In contrast to the control condition, mice selectively lick in anticipation of reward on go trials and specifically following the second odour, consistent with the use of relational structure. Importantly, when averaged across individual odours, overall licking is matched between control and structured tasks. **G.** Representative cFos staining in ventral CA1 (vCA1) following performance of the control and structured tasks. Scale bars = 2 mm, 300 µm, 100 µm. **H.** Quantification of cFos expression in vCA1. Structured task performance is associated with increased neuronal activity relative to the control condition. **I–J.** As **in G–H**, for medial prefrontal cortex (mPFC). In contrast to vCA1, cFos expression is reduced in the structured task relative to control.

In the structured task, half of trials are rewarded and half are unrewarded. Each trial contains two odour presentations, and odours are fully counterbalanced across positions and outcomes. As a consequence, each individual odour is rewarded on 25% of its presentations. Our control task exploited this property by delivering reward probabilistically on each odour presentation with the same overall probability (25%), independent of cue identity, order, or trial structure. Under these conditions, sensory stimulation, reward frequency, and contingency were matched to the structured task, but no relational information could be used to predict outcome.

Mice trained on the structured task learned as described above, reaching stable expert performance, whereas mice trained on the control task remained at chance performance throughout training (Fig. 4A-F). Importantly, when licking behaviour was averaged across all presentations of odours A, B, and C, there was no difference between structured-task and control-task mice, indicating comparable levels of motivation, reward engagement, and motor output across cohorts. Thus, the two tasks differed specifically in whether relational structure was behaviourally informative.

To confirm that this distinction was reflected in the strategies used by the animals, we applied the same regression and reinforcement learning analyses described above (Fig.4 – Figure Supplement 1). Behaviour in the structured task was best explained by structural predictors and was associated with high structural scores in the reinforcement learning model. In contrast, behaviour in the control task was poorly explained by structural predictors and remained near zero on the structural–elemental index, indicating that mice did not adopt a structured representation when no such structure was available.

We next used these two tasks to examine brain-wide engagement associated with learning and using temporal structure. Where possible, control-task mice were yoked to structured-task mice, such that they experienced the same number of training sessions and equivalent task exposure. Once structured-task mice had reached stable expert performance, animals were perfused 90 minutes after the final session and brains were processed for cFos immunohistochemistry. We quantified cFos-positive nuclei per mm² in ventral CA1, medial prefrontal cortex, nucleus accumbens shell, anterior thalamus and dorsal CA1.

We found that cFos expression was elevated in ventral CA1 in mice trained on the structured task compared to control-task mice (Fig. 4G,H). In contrast, cFos expression was reduced in medial prefrontal cortex (Fig. 4I,J), indicating that engagement of temporal structure is associated not with a global increase in activity, but with a redistribution of recruitment across circuits. These large changes were accompanied by a subtle increase in nucleus accumbens shell and no detectable change in anterior thalamus, lateral accumbens shell, or dorsal CA1 (Fig. 4 – Figure Supplement 2), consistent with selective engagement of specific circuits under matched sensory and reward conditions.

These results indicate that learning and using temporal structure selectively recruits ventral hippocampal circuitry while reducing engagement of medial prefrontal cortex. Because the two tasks are matched for sensory input, reward statistics, and motor output, this dissociation is unlikely to reflect generic differences in arousal, reward expectation, or licking, and instead suggests that distinct circuit motifs are engaged depending on whether behaviour can be guided by simple value contingencies or requires relational structure.

Together, these findings demonstrate that learning and using temporal structure selectively engages ventral hippocampal circuitry, providing a direct link between relational structure in behaviour and hippocampal circuit recruitment.

## DISCUSSION

The ability to extract structure from past experience and use it to guide future behaviour is fundamental to adaptive cognition^1–3^. From navigating familiar environments to interpreting the temporal order of events, animals must learn relationships between distinct cues and maintain these representations across time in order to make accurate predictions^4,7,16,17,42,43^. Despite its centrality, the neural basis of such structural learning has been difficult to isolate experimentally, in part because many commonly used behavioural paradigms confound relational structure with cue identity, value, or sensory binding^12,13,44,45^.

In the present study, we introduce a temporally structured olfactory task for mice that isolates relational structure while controlling for sensory input and reward statistics, providing a minimal and experimentally tractable assay of structural learning. By presenting cues sequentially, counterbalancing odour identity and position, and equating reward probability across individual cues, the task requires animals to rely on the ordered relationship between cues rather than elemental or configural strategies^12,44–46^. Using this design, we show that mice rapidly acquire structure-guided behaviour and that learning proceeds through a gradual shift towards structural representations.

Critically, this dissociation allows us to demonstrate that identical sensory inputs are assigned distinct behavioural values depending solely on their position within a learned structure. Dopamine activity provides an independent readout of this process, showing that value-based signals differentiate identical cues according to their temporal context^33,47–49^. This indicates that value assignment during behaviour depends on relational structure rather than cue identity^3,15,33,49^.

Finally, using a carefully matched control task, we show that ventral hippocampus is selectively engaged when structure is behaviourally relevant, even when sensory input and reward statistics are held constant^12,37,50^. Because these tasks are matched for sensory, reward, and motor variables, this difference isolates relational structure itself as the critical driver of hippocampal recruitment.

Together, these results establish a minimal, non-spatial paradigm for isolating relational structure in behaviour, demonstrate that animals use this structure to guide value-based predictions, and identify hippocampal circuitry as selectively recruited when such representations are required.

### Relation to spatial and configural accounts of relational learning

Relational learning has historically been studied in the context of spatial navigation, where animals must learn relationships between landmarks, trajectories, or locations^3,5,7,16^. Spatial tasks have provided compelling evidence for relational representations and their dependence on hippocampal circuitry^3,28,51^. However, spatial structure is only one instance of relational organisation, and it is often difficult to dissociate relational encoding from specialised spatial mechanisms or from continuous sensory cues.

An alternative approach has been provided by configural learning theory, in which compound stimuli are treated as distinct configurations rather than as collections of independent elements^13,14^. While configural accounts can explain performance in many biconditional and paired-associate tasks, they are often difficult to distinguish from genuinely relational strategies, particularly when cues are presented simultaneously and can be perceptually bound^12,13,29,44,45^. More generally, even tasks based on sequential cues do not necessarily isolate relational structure. In many cases, performance can be supported by attending selectively to one privileged part of the sequence, such as the first or last event, or by learning a coarse representation of the overall cue combination, without encoding the ordered relationship between overlapping elements. As a result, many tasks nominally described as “structural” can in practice be solved without explicit representation of relationships between discrete events^13,52^.

The task introduced here addresses this ambiguity directly. By separating cues across time and designing reinforcement contingencies such that no individual cue, unordered pair, or fixed sequence position is predictive, the task cannot be solved by perceptual binding or configural representations^12,44–46^. Instead, successful performance requires encoding the temporal order of cues as a relational structure^12,42,43,53^. In this sense, the task more closely parallels classical structural learning paradigms in humans, in which patients with hippocampal damage show impairments despite intact elemental learning^54–57^.

### Temporal structure as a model for episodic-like representations

A defining feature of episodic memory is the ability to represent sequences of events extended over time and to use recent past information to disambiguate the present^6,58^. When the same cue appears in different temporal contexts, accurate prediction requires combining information about what is happening now with what has just occurred, often referred to as the use of temporal context^4,28,42,43^. This process allows the construction of distinct internal “states” for otherwise identical sensory events^5,9,15,47,51^.

The present task captures this requirement in a minimal and experimentally tractable form. Identical odours evoke different behavioural and neural responses depending solely on their position within a sequence. Importantly, this distinction is expressed not only in behaviour but also in dopamine prediction errors, which reorganise across learning to reflect inferred task structure rather than sensory identity. This finding is consistent with the idea that dopamine signals reflect value estimates defined by internally constructed states, rather than being hard-wired to specific stimuli or actions^33^.

### Neural representations of structure: dopamine and hippocampus

Our results identify two complementary neural signatures of structural learning. First, dopamine activity in the nucleus accumbens reflects prediction errors aligned with structured representations, with anticipatory signals emerging selectively on rewarded sequences and outcome-related errors diminishing as learning progresses. Trial-by-trial and animal-to-animal variability in dopamine dynamics is captured by the same reinforcement learning models that describe behavioural strategy, indicating a tight link between representation, learning, and neuromodulatory signalling^33,48^.

Second, we find that learning a structured task selectively engages ventral CA1, as revealed by increased cFos expression compared to a carefully matched control task^50,59,60^. Because sensory input, reward statistics, and motor output were equated across tasks, this differential recruitment cannot be attributed to general arousal, motivation, or reward expectation. Instead, it is consistent with a role for hippocampal circuitry in representing and selectively engaging with relational structure during behaviour^12,42,53^. Interestingly, we did not observe a corresponding change in dorsal CA1 cFos, a region previously implicated in spatial forms of structural learning^37^. This suggests that temporal and spatial structure may preferentially engage distinct hippocampal subregions.

An additional feature of the cFos data is that the structured task was associated not only with increased activity in ventral CA1, but also with reduced activity in medial prefrontal cortex relative to the matched control task. Although this observation is correlational, it fits with the idea that these regions make dissociable contributions to behaviour^3,47,61^. In the control task, where reward is governed by simple probabilistic contingencies and no relational structure can be exploited, behaviour may rely more strongly on circuits involved in value-based prediction. By contrast, when successful behaviour depends on disambiguating overlapping cue sequences across time, recruitment shifts towards hippocampal circuitry capable of representing relational structure. In this view, hippocampus contributes the structured state representation on which downstream value signals, including dopamine prediction errors, can then operate^3,27,33,62^.

Together, these findings support a framework in which hippocampal circuits contribute to the construction or maintenance of structured representations, which are then used by downstream systems to compute prediction errors and guide behaviour^1,23^. While the present data do not establish a causal role for hippocampus, they provide a principled basis for future interventions that selectively target acquisition or expression of structure.

### Implications for disease models and future directions

Deficits in relational and episodic-like memory are hallmarks of a range of neuropsychiatric and neurodegenerative disorders, including Alzheimer’s disease, schizophrenia, and major depression^8–10^. Progress in understanding the cellular and circuit mechanisms underlying these impairments has been hindered by the lack of robust, high-throughput behavioural paradigms that isolate relational structure while remaining compatible with modern systems neuroscience approaches^12,13^.

The task introduced here offers several advantages in this regard. It is rapidly acquired, amenable to head-fixed recordings and manipulations, and designed such that structure can be selectively removed while preserving sensory and reward statistics. This makes it well suited for dissecting disease-related deficits in acquisition, generalisation, or expression of structured behaviour^8,9^. Moreover, the ability to quantify learning strategy continuously using regression and reinforcement learning models provides a powerful framework for linking behavioural phenotypes to neural mechanisms.

A key open question raised by these findings is how relational structure is represented and used to guide behaviour at the circuit level. Recent computational work provides a concrete framework for this^61^. In particular, successor representation models that incorporate feature–outcome structure can support inference over relationships between sequential cues by learning predictive maps of state transitions. In this formulation, value is assigned not to individual stimuli, but to their position within a learned structure of expected future states.

This framework offers a mechanistic account of how hippocampal circuits could support the transition from cue-based to structure-based strategies observed in our data. By encoding relationships between cues in a predictive map, hippocampal output could provide the inferred state representation required for downstream systems, including dopaminergic circuits, to compute prediction errors over inferred rather than directly observed states^3,27,33^. Consistent with this account, dopamine signals in our task reflect outcome predictions that depend on the sequential relationship between cues, rather than their individual identity.

Future work can exploit the flexibility of the task design to probe how structural representations are learned and expressed under different conditions. For example, varying the number of cues that change during generalisation, manipulating transition probabilities between cue pairs, or altering reward contingencies independently of structure would allow further dissociation of structure and value. Similarly, temporally precise perturbations during learning or performance could be used to dissect the contribution of distinct circuits at different stages of the task.

In summary, this study establishes a minimal, non-spatial paradigm for studying structural learning in mice, demonstrates that behaviour, dopamine signalling, and hippocampal engagement all reflect learned task structure, and provides a foundation for future mechanistic investigations of relational memory in health and disease.

## ACKNOWLEDGEMENTS

We thank members of the MacAskill laboratory for comments on the manuscript, and the biological services central unit at University College London for animal care and technical assistance. AFM is funded by a UKRI Frontier Research Fellowship EP/Y034724/1 and a Medical Research Council project grant number MR/W02005X/1. SN was funded via the UCL 4-year Sainsbury Wellcome Centre PhD, DO and MC were funded by the UCL/Birkbeck MRC Doctoral Training Program.

## AI STATEMENT

Artificial intelligence (ChatGPT, OpenAI) was used to assist with text and code editing. All scientific content, analyses, and interpretations were generated and verified by the authors.

## MATERIALS AND METHODS

### Animals

Adult male and female C57BL/6J mice aged 9 to 11 weeks at the start of experiments were obtained from Charles River. Mice were housed in groups of 1 to 4 under a 12 h light-dark cycle with food available ad libitum. Water access was restricted during behavioural training as described below. All procedures were carried out in accordance with UK Home Office regulations and University College London guidelines.

For the experiments reported here, mice were used for behavioural testing following implantation of a lightweight metal headbar. Animals underwent stereotaxic surgery and were allowed to recover for at least 7 days before behavioural procedures began.

### Stereotaxic surgery

Mice aged 7 to 12 weeks were anaesthetised with isoflurane in oxygen and placed in a stereotaxic frame on a feedback-controlled heating pad maintained at 35 to 37°C. Anaesthesia was induced at 4% isoflurane and maintained at 1 to 2% during surgery. Eyes were protected with ophthalmic gel. After scalp sterilisation and incision, the skull was cleaned and levelled using bregma and lambda. Local anaesthetic was applied to the scalp before craniotomy.

For experiments involving viral injection or implantation, small craniotomies were made at the required stereotaxic coordinates, and injections were delivered with a Nanoject II through pulled glass pipettes. Viral injections consisted of 250 to 500 nL delivered in 14 or 28 nL steps at 15 s intervals. After infusion, the pipette was left in place for 5 min before withdrawal. Optical fibres were implanted as required and secured with dental cement. Injection sites were verified histologically at the end of experiments. All mice received carprofen during surgery and for 48 h post-operatively. Animals were monitored during recovery and returned to their home cage only once fully ambulatory.

### Histology

For histological verification, mice were deeply anaesthetised with ketamine and xylazine and transcardially perfused with ice-cold 4% paraformaldehyde. Brains were dissected, post-fixed overnight at 4°C, and transferred to phosphate-buffered saline. Viral expression and implant placement were verified using serial-section two-photon imaging of whole brains.

### Behavioural apparatus

Mice were trained in a head-fixed olfactory paired-associates task while running on a cylindrical treadmill. The treadmill consisted of a 3D-printed wheel suspended on a metal axle, allowing one-dimensional locomotion^63^. Headbars were secured to a custom holder with adjustable height and angle^64^.

Odours were delivered through a custom olfactometer adapted from previously described designs^40^. A constant stream of clean carrier air was directed towards the animal, while a vacuum line continuously removed air from the opposite side of the snout to minimise residual odour contamination. Odour stimuli were generated by routing pressurised air through one of several glass vials containing pure odorants and injecting the resulting odourised air into the carrier stream. Odour valves were opened for 1 s, after which airflow returned to clean air only. Carrier and odourised airflows were matched at approximately 0.7 L/min.

Odours were selected from a panel of neutral odorants previously shown not to be inherently appetitive or aversive to mice and were chosen to maximise perceptual separability. Odour delivery was characterised using a mini photoionisation detector placed at the position of the mouse snout. Because detector amplitude cannot be directly compared across different odorants, airflow was also monitored with an airflow sensor to confirm matched delivery across cues.

Licking was measured using a capacitive lick detector (AT42QT1011, SparkFun) connected to a metal lick spout positioned in front of the animal. Water rewards of approximately 10 µL were delivered through the lick spout by solenoid valve. Behavioural control was implemented using custom Arduino-based software, with event timing and analogue lick signals recorded through a National Instruments data acquisition board.

### Behavioural training

After at least 7 days of post-operative recovery, mice were water restricted to approximately 85% of their baseline body weight. Following at least 1 week of handling and water restriction, mice were habituated to head fixation over 4 days, first on a static platform and then on the treadmill. On the fifth day, mice received water from the lick spout at pseudorandom intervals to habituate them to reward collection under head fixation.

Mice were then trained on an olfactory paired-associates task. Each trial consisted of two sequential odour presentations separated by a delay. In the initial shaping phase, only rewarded odour pairs were presented, and water was delivered 1.5 s after the second odour regardless of behaviour. Mice advanced to the full task once they showed anticipatory licking during the response window on at least 70% of shaping trials.

In the full task, rewarded and unrewarded odour pairs were presented pseudorandomly, with all trial types represented within each block. Rewarded trials were followed by water delivery 1.5 s after the second odour. Unrewarded trials were not punished, but no reward was delivered. Mice typically learned to withhold licking on unrewarded trials within 8 to 10 days. Behaviour was scored based on licking during the response window after the second odour. Trials were classified as hits, misses, false alarms, or correct rejections according to trial type and licking behaviour. A typical session consisted of approximately 80 trials. Daily water intake was maintained at 1 to 1.5 mL, delivered during or after the session.

### Behavioural control experiments

Several control conditions were used to exclude non-olfactory or configural explanations for performance. In auditory-only control sessions, airflow was turned off and the odour delivery tube was disconnected, such that the only available cues were valve clicks. In sensory-plus-auditory control sessions, carrier airflow remained on but odour vials were replaced with empty vials, preserving valve sounds and airflow changes in the absence of odours. The system was flushed for at least 30 min before such sessions.

To test whether animals could have solved the task using residual odour mixtures rather than sequential structure, a diluted version of one odour was delivered continuously throughout the trial on a subset of trials, followed by the second odour. This manipulation created an explicit low-concentration mixture signal. If mice relied on a configural mixture cue formed by overlap between the first and second odours, performance should have remained high on these trials.

### Behavioural analysis

Behavioural performance was quantified as the proportion of correct trials, defined as the sum of hits and correct rejections divided by the total number of trials, or as sensitivity, quantified using d-prime, calculated as the difference between the z-transformed hit rate and false alarm rate.

### Logistic regression analysis

To quantify the contribution of different task features to behaviour, we used logistic regression models to predict anticipatory licking on each trial from task variables. Predictor sets included elemental odour identity, unordered odour configurations, ordered odour pairs, trial history variables, and a bias term. Models were fit using scikit-learn. Data were split into training and test sets, hyperparameters were optimised by cross-validation, and model performance was evaluated on held-out data. To estimate the contribution of specific groups of predictors, we compared both the performance of each predictor alone, and also the performance of the full model with reduced models lacking the predictors of interest. This allowed us to quantify the relative influence of elemental, structural, and history-based information on behaviour.

### Reinforcement learning models

To characterise how behaviour evolved over learning, we fit a family of Rescorla–Wagner reinforcement learning models adapted from ref^29^ to trial-by-trial licking behaviour. These models differed in whether they learned values for individual odour elements, ordered odour pairs (structural model), or both (combined model). This allowed us to compare behavioural strategies based on elemental learning, structural learning, and their combination.

For each trial, the behavioural dataset comprised the first odour, second odour, trial outcome, licking response, and ordered trial type. Licking was treated as a binary choice variable, defined as 1 when the animal produced an anticipatory lick and 0 otherwise.

#### Model structure

We fit three model classes independently to each mouse and training day. The elemental model learned values for individual odours. On each trial, the predicted tendency to lick was determined by the weighted sum of the learned values associated with the first and second odours. The structural model learned values for ordered odour pairs. In this model, each distinct sequence of first and second odours was assigned its own value. The combined model incorporated both representations simultaneously. On each trial, the decision variable was given by the sum of a structural component, an elemental component, and a constant bias term. The structural component corresponded to the value of the ordered odour pair multiplied by a pair-specific weight. The elemental component corresponded to the weighted sum of the values of the two constituent odours, allowing different contributions from the first and second odours. The contribution of the first odour was further scaled by a parameter 𝛾, enabling asymmetric weighting across the sequence. Choice probability was then obtained by passing the total decision variable through a logistic function.

Formally, on trial t, for odours 𝑜1 and 𝑜2, the probability of licking was:

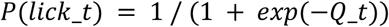

Where:

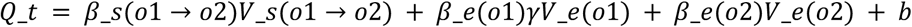

Here, 𝑉_𝑠 denotes the learned value of the ordered odour pair, 𝑉_𝑒 denotes the learned value of each individual odour, 𝛽_𝑠 and β_𝑒 are structural and elemental weighting parameters respectively, 𝛾 scales the contribution of the first odour within the elemental pathway, and 𝑏 is a constant bias term.

#### Learning rule

Value updates followed a standard Rescorla-Wagner rule. For the structural component, the value of the experienced ordered odour pair was updated according to:

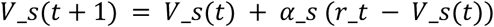

where 𝑟_𝑡 was the reward delivered on trial t, and 𝛼_𝑠 was the structural learning rate.

For the elemental component, the values of both odours presented on that trial were updated independently according to:

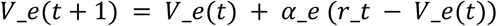

where 𝛼_𝑒 was the elemental learning rate. All values were initialised to zero at the start of each fitting run.

Thus, in the combined model, ordered odour-pair values and individual odour values were learned in parallel from the same reward signal and jointly influenced choice.

#### Model fitting

Models were fit separately for each mouse and day. Model parameters were estimated by maximising the likelihood of the observed sequence of licking responses given the trial-by-trial task history.

For a given parameter set, the model was run sequentially through the session. On each trial, the model generated a predicted lick probability before updating its internal values from the observed reward outcome. The log-likelihood of the observed licking data was then computed under a Bernoulli observation model:

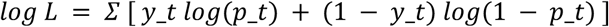

where 𝑦_𝑡 is the observed lick response on trial t, and 𝑝_𝑡 is the model-predicted probability of licking. Probabilities were clipped to the interval [10^-6^, 1-10^-6^] to avoid numerical instability.

Parameters were estimated by minimising the negative log-likelihood using the L-BFGS-B algorithm implemented in SciPy. To reduce sensitivity to initialisation, each model was fit from multiple random starting points, and the solution with the lowest negative log-likelihood was retained. Parameter bounds were constrained as follows: learning rates were bounded between 0.01 and 1, inverse temperature terms between 0.1 and 10, the elemental scaling term gamma between 0 and 1, and the bias term between -5 and 5.

#### Model comparison

We compared model variants using Akaike Information Criterion (AIC), calculated from the maximised log-likelihood and the number of free parameters in each model. The number of parameters depended on model class. The structural-only model contained a structural learning rate, one structural weight per ordered odour pair, and a bias term. The elemental-only model contained an elemental learning rate, one elemental weight per odour, the scaling parameter, and a bias term. The combined model contained both sets of parameters.

These comparisons allowed us to determine whether behaviour was better captured by learning about individual odours, ordered odour pairs, or a combination of both.

#### Learning style score

To summarise the relative contribution of structural and elemental learning at the session level, we computed a learning style score from the fitted likelihoods of the three model classes. This metric quantified whether the combined model was better approximated by the structural-only or elemental-only model, with positive values indicating relatively greater structural influence and negative values indicating relatively greater elemental influence.

### Dopamine photometry

#### Viral expression and fibre implantation

For photometry experiments, mice were injected with 200–400 nL of AAV5-CAG-dLight1.1 into the nucleus accumbens (NAc) to enable optical measurement of dopamine release. A fibre optic cannula (200 µm core diameter, 0.39 NA, 5 mm length; Thorlabs) was implanted unilaterally above NAc during the same surgery. Implants were secured to the skull using dental cement anchored to skull screws, and the surrounding skin was sealed with surgical adhesive.

#### Photometry acquisition

Dopamine-dependent fluorescence was recorded using a fibre photometry system as described previously^27,65,66^. Excitation light at 470 nm was used to drive dLight1.1 fluorescence, while a 405 nm channel was used to capture dopamine-independent fluctuations in signal.

To allow separation of the two signals, excitation light from the 470 nm and 405 nm LEDs was sinusoidally modulated at distinct frequencies (500 Hz and 210 Hz, respectively). Signals were combined optically and delivered via a fibre patch cord to the implanted cannula. Emitted fluorescence was collected through the same fibre, filtered (emission >505 nm), and detected using a photoreceiver sampled at 10 kHz.

Signals corresponding to the two excitation wavelengths were separated offline by demodulation. Recordings were synchronised to behavioural events via TTL pulses sent from the behavioural control system at the start of each session.

#### Signal preprocessing

Photometry data were processed using custom Python scripts. Both 470 nm and 405 nm signals were low-pass filtered to reduce high-frequency noise. To correct for photobleaching, a fourth-order polynomial was fit to each trace and subtracted. Movement-related and other non-specific fluctuations were estimated by regressing the 405 nm signal onto the 470 nm signal using a least-squares linear fit. The fitted component was subtracted from the 470 nm trace to yield a movement-corrected fluorescence signal corresponding to dopamine dynamics. The corrected signal was then z-scored within session.

#### Event alignment and summary measures

To quantify dopamine responses, signals were aligned to task events on a trial-by-trial basis. Two key epochs were analysed: a predictive period following second odour presentation, and a post-outcome period following reward delivery or omission.

For each trial, the signal was baseline-corrected using the mean fluorescence in the 1 s preceding the first odour. Event-related responses were quantified as the mean z-scored fluorescence within 1s time windows following each event. These windows were chosen to capture the majority of the evoked signal while remaining robust to variability in response timing across animals.

#### Regression analysis of dopamine signals

To relate dopamine signals to task variables, we applied a regression framework analogous to that used for behaviour. For each trial, dopamine activity within the predictive and post-outcome windows was used as the dependent variable in a linear regression model. Predictors included elemental odour identity, ordered odour pairs (structural predictors), and trial history variables. Models were fit using ordinary least squares, and predictor weights were used to quantify the contribution of each feature to dopamine signals. This approach allowed us to directly compare how behavioural output and dopamine activity were shaped by elemental and structural information, and how these representations evolved over learning.

#### Relationship between dopamine signals and learning strategy

To relate dopamine signals to behavioural learning strategy, we leveraged the reinforcement learning model fits described above. For each session, we computed a learning style score, which quantified the relative contribution of structural versus elemental learning based on model likelihoods. Dopamine responses were then summarised at the session level. For each session, we computed the mean z-scored dopamine signal across trials within predefined task epochs (predictive and post-outcome), restricting analysis to rewarded (go) trials. These trials provide a consistent context in which cue-evoked predictions are behaviourally relevant and comparable across sessions.

Session-averaged dopamine responses were then related to the corresponding learning style score. Each session therefore contributed a single datapoint relating dopamine activity to inferred learning strategy. This analysis allowed us to test whether sessions in which behaviour was more strongly driven by structural versus elemental learning were associated with systematic differences in predictive dopamine signalling.

### c-Fos immunohistochemistry

#### Tissue collection and staining

To assess neuronal activity across experimental conditions, mice were transcardially perfused 90 minutes after the start of the behavioural session. This timepoint was chosen to capture activity-dependent c-Fos expression associated with task performance. Animals were perfused with 20 mL of ice-cold phosphate-buffered saline (PBS), followed by 20 mL of ice-cold 4% paraformaldehyde (PFA).

Brains were extracted, post-fixed, and sectioned. Free-floating sections were washed in PBS and incubated in a blocking solution containing 2% bovine serum albumin and 0.5% Triton X-100 in PBS to reduce non-specific binding. Sections were then incubated overnight at 4°C with a primary antibody against c-Fos (mouse monoclonal, 1:500) diluted in blocking solution.

The following day, sections were washed in PBS and incubated for 1 hour at room temperature with a fluorescent secondary antibody (Alexa Fluor 555 donkey anti-mouse, 1:1000). After three additional washes in PBS, sections were mounted onto glass slides with DAPI-containing mounting medium, coverslipped, and sealed prior to imaging.

Coverslipped sections were imaged using a Zeiss Axio Scan slide scanner. Imaging parameters, including exposure time and illumination intensity, were optimised for each fluorophore and held constant across samples within each experiment to ensure comparability.

#### Data processing and quantification

Cell detection was performed using QuPath with watershed-based automated cell detection. Detection parameters were manually optimised to ensure consistent identification of labelled nuclei across sections. Cell detection outputs, including spatial coordinates and fluorescence intensity, were processed using a custom Python pipeline.

To exclude non-somatic signal sources such as debris or antibody aggregates, detections were filtered based on nuclear circularity. Only objects with a circularity score ≥ 0.45 were retained for analysis. For each region and animal, we quantified the density of c-Fos-positive cells.

### Statistical analysis

Statistical analyses were performed in Python using scipy, statsmodels and pingouin. Summary data are reported as mean ± s.e.m. Normality was assessed by visual inspection of the data. Where appropriate, to account for variability across animals, we used linear mixed-effects models with experimental condition as a fixed effect and animal identity as a random intercept. Statistical details are provided in the main text and figure legends. The threshold for significance was p < 0.05. Sample sizes were chosen on the basis of previous studies, and no formal power calculations were performed.

## SUPPLEMENTARY STATISTICS TABLE

| Figure | Description | n | Test | Statistic | p-value |
| --- | --- | --- | --- | --- | --- |
| 1F | p-correct x day, per mouse | 14 | Repeated measures ANOVA | $F(9,45) = 11.54$ | <0.001 |
| 1G | p_correct day 1 vs day 9 | 14 | Paired T-test | $t(13) = 10.83$ | <0.001 |
| | d prime day 1 vs day 9 | 14 | Paired T-test | $t(13) = 8.34$ | <0.001 |
| 2C | Single predictor $R^2$ x predictor class | 14 | Repeated measures ANOVA | $F(3,42) = 41.40$ | <0.001 |
| 2D | Delta $R^2$ (unique variance) x predictor class | 14 | Repeated measures ANOVA | $F(3,42) = 41.57$ | <0.001 |
| 2F | $\Delta AIC$ x day x model type | 14 | 2-way repeated measures ANOVA | $F(9,54) = 3.33$ (day)<br>$F(1,6) = 1.04$ (model)<br>$F(9,54) = 5.42$ (interact) | 0.047<br>0.347<br>0.016 |
| | DAIC structure vs elemental day 1 | | Paired T-test | $t(13) = 3.71$ | 0.002 |
| | DAIC structure vs elemental day 9 | | Paired T-test | $t(11) = 3.64$ | 0.004 |
| | Learning style score x day | | Repeated measures ANOVA | $F(9,54) = 5.06$ | <0.001 |
| 3B | Preemptive dopamine x day x go/nogo | 6 | 2-way repeated measures ANOVA | $F(1,5) = 0.85$ (day)<br>$F(1,5) = 86.36$ (go/nogo)<br>$F(1,5) = 77.17$ (interact) | 0.44<br><0.001<br><0.001 |
| | Outcome dopamine x day x go/nogo | | | $F(1,5) = 1.97$ (day)<br>$F(1,5) = 7.41$ (go/nogo)<br>$F(1,5) = 10.94$ (interact) | 0.22<br>0.041<br>0.021 |
| 3D | Single predictor $R^2$ x model x stage | 6 | 2-way repeated measures ANOVA | $F(1,5) = 23.36$ (stage)<br>$F(3,15) = 34.58$ (model)<br>$F(3,15) = 33.82$ (interact) | 0.005<br><0.001<br><0.001 |
| 3E | $\Delta R^2$ (unique variance) x model x stage | 6 | 2-way repeated measures ANOVA | $F(1,5) = 24.38$ (stage)<br>$F(3,15) = 30.69$ (model)<br>$F(3,15) = 32.77$ (interact) | 0.0043<br>0.0015<br>0.0017 |
| 3G | Learning style score x preemptive dopamine (go trials) | 6 mice, 83 sessions | Linear mixed-effects model (random intercept: mouse) | $\beta = 0.523$<br>$z = 6.02$ | <0.001 |
| | Learning style score x outcome dopamine (go trials) | | | $\beta = -0.172$<br>$z = -2.67$ | 0.008 |
| 4C,F | p_lick x group x go/nogo | 7 mice (3 control, 4 structure) | Mixed ANOVA | $F(1,5) = 0.07$ (group)<br>$F(1,5) = 29.95$ (go/nogo)<br>$F(1,5) = 15.31$ (interact) | 0.802<br>0.003<br>0.011 |
| | Control (go vs nogo) | 3 mice | t-test | $t(2) = 1.60$ | 0.250 |
| | Structure (go vs nogo) | 4 mice | t-test | $t(3) = 5.35$ | 0.013 |
|  | p_lick × group × odour | 7 mice (3 control, 4 structure) | Mixed ANOVA | F(1,5) = 0.07 (group)<br>F(2,10) = 0.11 (odour)<br>F(2,10) = 0.19 (interact) | 0.804<br>0.893<br>0.830 |
| 4F |  |  |  |  |  |
| 4H | vCA1 cFos density × task (control vs structured) | 7 mice, 14 samples | Linear mixed-effects model (random intercept: mouse) | $\beta = 0.891$<br>$z = 3.46$ | 0.001 |
| 4J | mPFC cFos density × task (control vs structured) | 6 mice, 12 samples | Linear mixed-effects model (random intercept: mouse) | $\beta = -0.632$<br>$z = -2.26$ | 0.024 |
| 1 Sup1A | Performance × airflow condition | 5 mice | Repeated measures ANOVA | F(2,8) = 33.02 | < 0.001 |
| 1 Sup1A | Performance × odour condition | 4 mice | Repeated measures ANOVA | F(2,6) = 9.77 | 0.013 |
| 1 Sup1I | p_lick × odour identity (A, C, C_low) | 3 mice | Repeated measures ANOVA | F(2,4) = 39.41 | 0.002 |
|  | Pairwise comparisons<br>A vs C<br>A vs C_low<br>C vs C_low |  | Planned paired t-test | t(2) = -0.65<br>t(2) = 5.29<br>t(2) = 10.66 | 0.585<br>0.034<br>0.009 |
| 3 Sup1B | Odour A (go vs nogo) | 7 mice | t-test | t(5) = 7.09 | <0.001 |
|  | Odour B (go vs nogo) |  | t-test | t(5) = 2.64 | 0.046 |
|  | Odour C (go vs nogo) |  | t-test | t(5) = 3.96 | 0.011 |
| 4 Sup1B | Single predictor R<sup>2</sup> × model × condition | 7 mice (3 control, 4 structure) | Mixed ANOVA | F(1,5) = 33.34 (condition)<br>F(3,15) = 24.10 (model)<br>F(3,15) = 24.12 (interact) | 0.002<br><0.001<br><0.001 |
| 4 Sup1C | $\Delta R^2$ (unique variance) × model × condition | 7 mice (3 control, 4 structure) | Mixed ANOVA | F(1,5) = 30.04 (condition)<br>F(3,15) = 26.93 (model)<br>F(3,15) = 20.65 (interact) | 0.003<br><0.001<br><0.001 |
| 4 Sup1E | Learning style score × condition | 7 mice (3 control, 4 structure) | Welch's t-test | t(8.53) = 6.00 | <0.001 |
| 4 Sup2 | Medial NAc cFos density × task (control vs structured) | 6 mice, 12 samples | Linear mixed-effects model (random intercept: mouse) | $\beta = 0.514$<br>$z = 1.99$ | 0.046 |
| | Lateral NAc cFos density × task (control vs structured) | | | $\beta = 0.168$<br>$z = 0.810$ | 0.418 |
| | Anterior Thalamus cFos density × task (control vs structured) | | | $\beta = 0.091$<br>$z = 0.422$ | 0.673 |
| | dCA1 cFos density × task (control vs structured) | | | $\beta = 0.051$<br>$z = 0.318$ | 0.750 |

---

**Figure 1-figure supplement 1.**
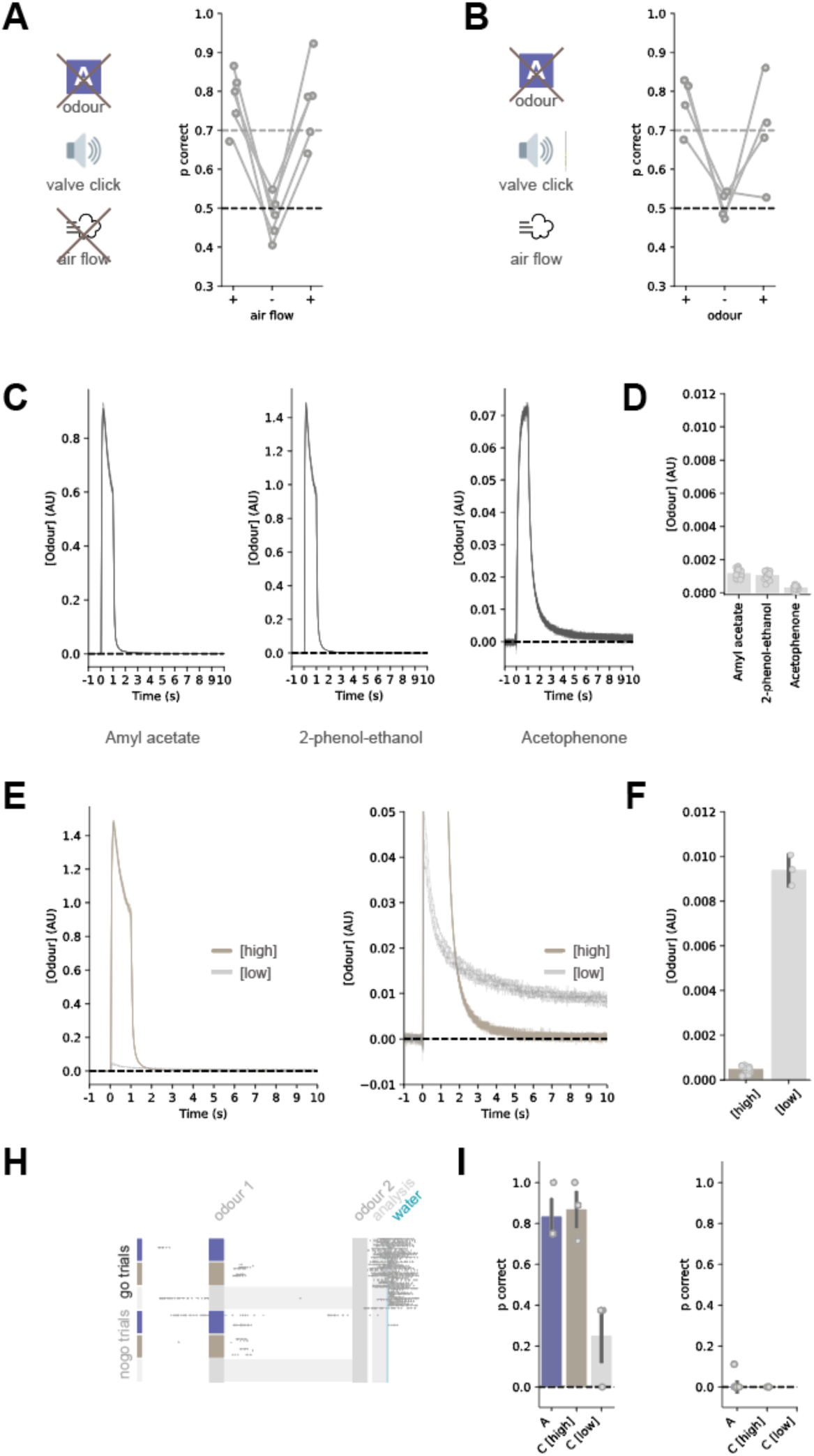
Control experiments exclude simple sensory explanations of performance. **A.** Removal of both odour delivery and airflow reduces task performance to chance, confirming that behaviour depends on olfactory input. **B.** Removal of odour while maintaining neutral airflow also reduces performance to chance, ruling out the use of airflow or other non-olfactory cues. **C.** Photoionisation detector (PID) measurements showing reliable odour delivery with rapid return to baseline following stimulus offset. **D.** Quantification of baseline odour levels at the time of second odour presentation, indicating minimal residual odour under standard conditions. **E.** Calibration of a control odour dilution designed to exceed residual odour levels, used to test whether animals could use lingering odour to solve the task. **F.** PID measurements during probe trials showing elevated residual odour concentrations prior to second odour delivery under the dilution manipulation. **G.** Example lick rasters from probe trials with elevated residual odour. Despite increased background odour levels, mice do not show anticipatory licking, indicating that residual odour is insufficient to guide behaviour. **H.** Left, proportion correct for ‘go’ pairs starting with odour A (unmanipulated), odour C (baseline), and odour C with added diluted odour. Performance is markedly reduced in the dilution condition. Right, performance on no-go trials is unchanged, consistent with a floor effect.

**Figure 2-figure supplement 1.**
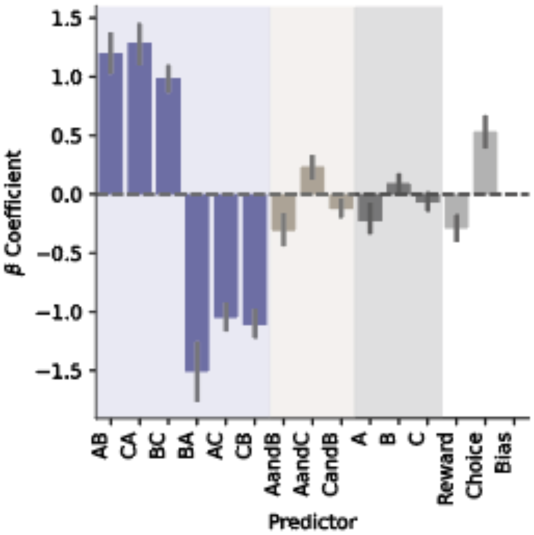
Regression coefficients selectively capture behaviourally relevant cue pairs. Regression coefficients (β weights) from the behavioural model (Figure 2). Task-relevant cue pairs show large coefficients with appropriate sign (positive for rewarded pairs, negative for unrewarded pairs), whereas coefficients for individual odours and other predictors remain small. This indicates that behaviour is primarily driven by structured cue combinations rather than elemental features or nonspecific correlations.

**Figure 3-figure supplement 1.**
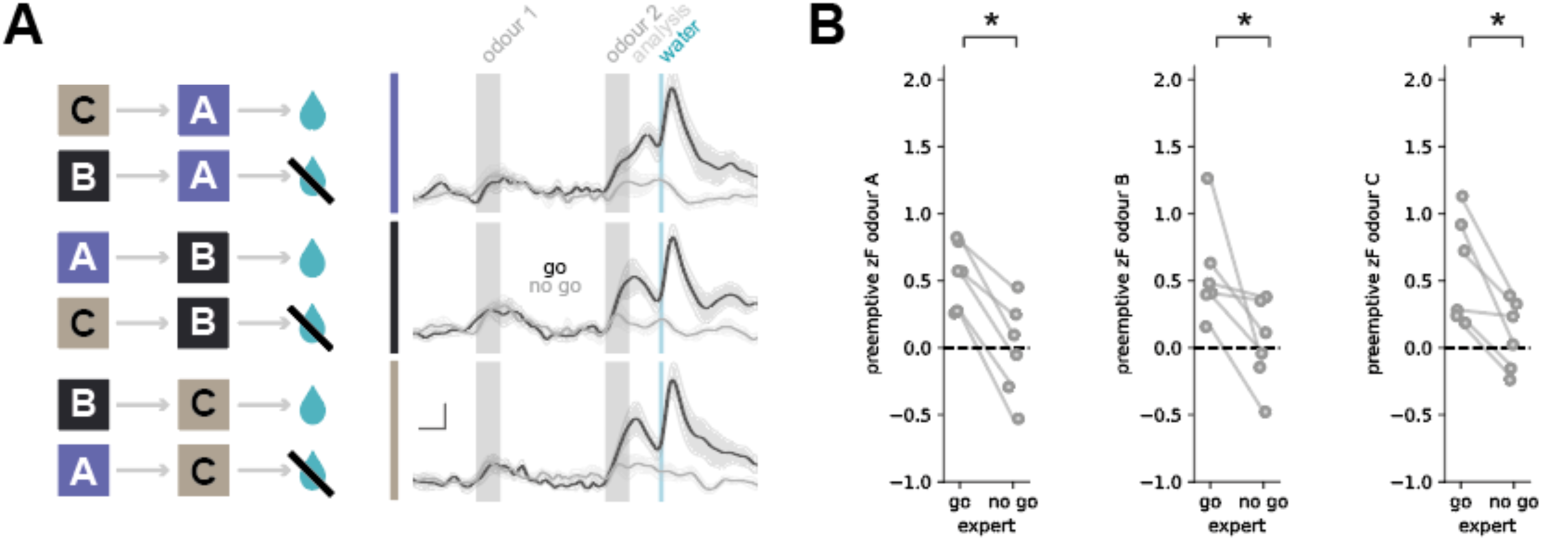
Differential dopamine responses to identical sensory stimuli dependent on temporal context. **A.** Dopamine traces for go and no-go trials, separated by the identity of the second odour. Clear separation of predictive dopamine signals is observed despite identical sensory input. Scale bar: 1 s, 0.5 zF. **B.** Quantification of pre-emptive dopamine responses for each second odour.

**Figure 4-figure supplement 1.**
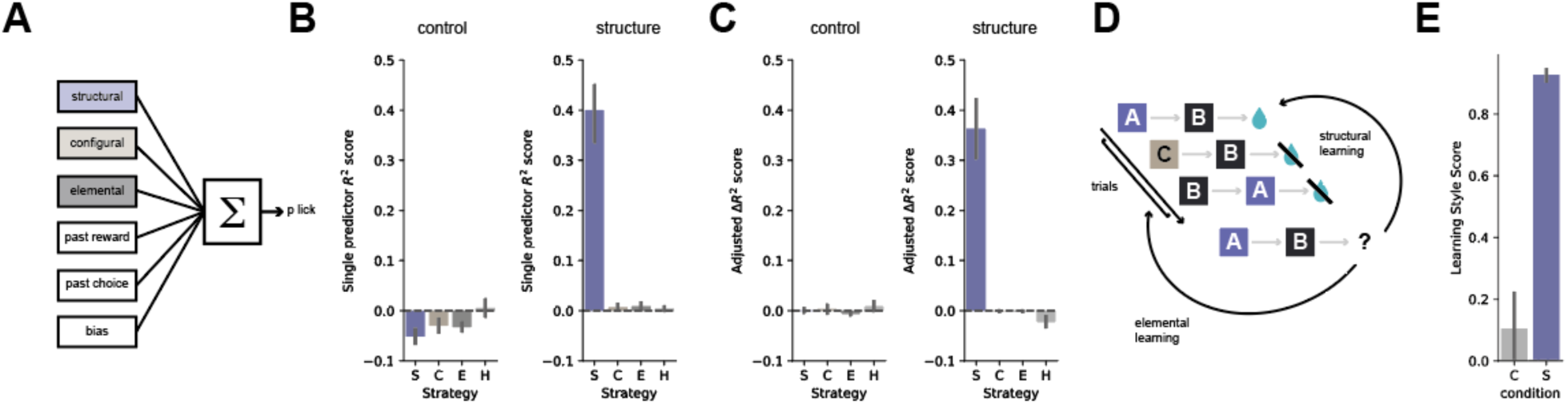
Behaviour of cFos cohort is well captured by structure-based models. A. Regression framework used to predict trial-by-trial licking behaviour, as in Figure 2, including structural, configural, elemental, and history predictors (see Methods). B. Variance explained by individual predictor classes (single predictor pseudo-R²) for control and structured tasks. In the control task, all predictors explain minimal variance, whereas in the structured task, structural predictors account for the majority of explained variance, with configural, elemental, and history predictors contributing little. C. Unique variance explained (Δpseudo-R²) by each predictor class for control and structured tasks in the same cohort. As in B, predictors contribute minimally in the control task, while in the structured task the majority of unique variance is captured by structural predictors, indicating adoption of a structure-based strategy. D. Schematic of the reinforcement learning models used to quantify behavioural strategies. E. Learning style score (difference between structural and elemental model fits) for control and structured tasks. Scores are near zero in the control task and strongly positive in the structured task, confirming selective engagement of structure-based learning in the cFos cohort.

**Figure 4-figure supplement 2.**
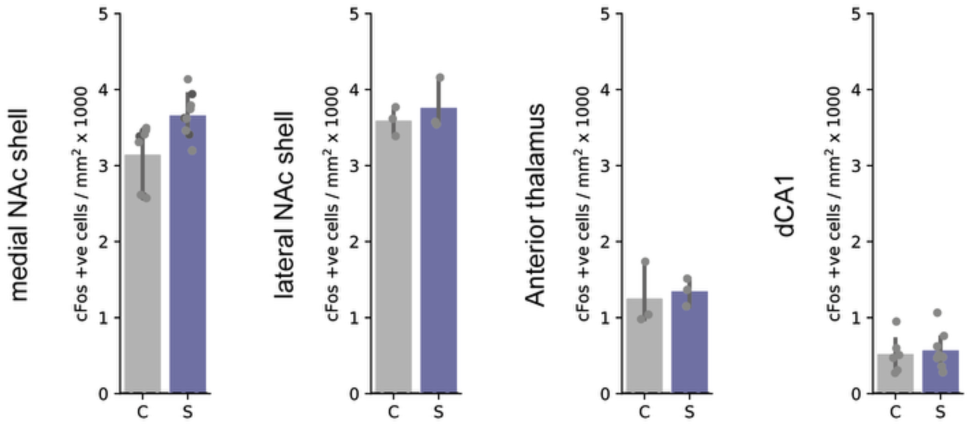
cFos changes are region-selective and extend to a downstream ventral striatal target. Quantification of cFos expression in additional brain regions following performance of the control and structured tasks. Left, medial nucleus accumbens (NAc) shell; middle, lateral NAc shell, anterior thalamus, right dorsal CA1. Medial NAc shell shows a significant but smaller increase in cFos expression in the structured task, whereas lateral NAc shell, anterior thalamus and dCA1 do not differ between conditions. These data indicate that structure-guided behaviour does not produce a global increase in activity, but instead recruits a selective circuit, with the largest effect observed in ventral CA1 and a smaller increase in medial NAc shell, consistent with engagement of a known downstream target.

